# Assessing the Effectiveness of Mycorrhizal Inoculation on the Biochemical Composition of the Different Species of Echinacea

**DOI:** 10.1101/2021.07.09.451863

**Authors:** Asha Sharma, Ishan Saini, Bandi Arpitha Shankar

## Abstract

The information regarding the effect of the mycorrhizal inoculation on different Echinacea species is not available in detail. Therefore, here we determined the changes in the biochemical composition of echinacea as a result of mycorrhizal inoculation. This experiment was undertaken to assess the effect of the mycorrhizal association on biochemical properties of different echinacea species (*E*. *angustifolia,E. purpurea, E*. *pallida*). Here various echinacea species were inoculated with mycorrhiza to examine the species richness in different traits. The results established that biological traits (plant dry matter, chlorophyll content, carotenoid, N content, P content, K content) and physiological and mycorrhization characteristics (Root essential oil, ABTS Antioxidant, Ferric-Reducing Antioxidant, Total phenolic, AM Spore No., AM Root Colonization) both are higher under mycorrhizal association than the control plants of different echinacea species. *E. purpurea* showed greater results than the *E*. *angustifolia* and *E*. *pallida.* Among biochemical properties chlorophyll content, carotenoid and N, P, K were significantly higher under*E*. *Purpurea* than the *E. angustifolia and E*.*pallida.* Total dry matter was higher under *E. angustifolia* (49.23 g) and minimum dry matter was found under *E*. *pallida* (40.07 g). Physiological and mycorrhizal traits were significantly higher under *E*. *purpurea* than the other species. *E*. *purpurea* showed higher AM Spore No., AM Root Colonization 231.30, 78.70% respectively. Lowest physiological and mycorrhization characteristics were found under *E*. *pallida.* The result of mycorrhizal association was very effective for plant growth and increased bio-physicochemical properties than the control plants.

## INTRODUCTION

Echinacea, also known as purple coneflower, is a member of the Asteraceae family. Botanically, echinacea is a perennial, herbaceous plant-primarily existing in eastern North America. Echinacea has an extended historical past of frequent use for an extensive range of illnesses.^[1]^ Medical studies validate a lot of conventional applications. It is probably one of the essential plant species that has more herbal healing value.

The plant is used in standard cold, bronchitis, coughs, some inflammatory conditions, upper respiratory infections plus urinary tract infection.^[2]^ The oldest record of its medicinal uses dates back to the use by the North American Indians for managing the infections and wounds. Later, in the late 1800s, these echinacea formulations began being contemporary as remedies for the standard cold.^[3]^ Moreover, preparations also vary owing to variations within the plant part used, the method used for the biochemical compound extraction, the geographic location.^[4]^ Despite the variability among echinacea compounds, attempts have been made to standardize as well as to characterize the substance used in medical studies.^[5]^ The properties of phytochemical constituents of echinacea roots have been analyzed for several oxidative stresses, including various types of cancers.^[6]^ In recent years, echinacea items are among the best-selling medicinal health cures from the planet. The taxonomic category, identification and also phylogenetic connection of echinacea species were previously developed according to the morphological and phytochemical variation.^[7]^ There are lots of parallels in morphology, physiology and chromosome quantity among echinacea species.^[8]^ Additionally, many species might have various ingredients with different effects that could decrease the security and usefulness of echinacea products. Consequently, it’s imperative to identify the plant species employed for therapeutic purposes properly the amounts of these biochemicals can also be enhanced by metabolic engineering. ^[9–11]^

In comparison to mutually advantageous mycorrhizal interactions, several mycoheterotrophic vegetation (approximately 400 plant species from various place families, pteridophytes, such bryophytes, as well as angiosperms) depend on mycorrhizal fungi for their CO2resources.[12] These plants have dropped their photosynthetic abilities and also parasitize mycorrhizal fungi which are connected with neighbor autotrophic vegetation. Arbuscular mycorrhizal fungi colonize the origins of numerous agriculturally crucial meal and bioenergy plants and may perform as’ biofertilizers and bioprotectors’ in eco-sustainable agriculture. Due to the health-promoting benefits of echinacea, it is crucial to increase the bioactive ingredients of the Echinacea by undertaking breeding and other conventional approaches ^[13]^. The significant variations present in caffeic acid as well as alamedas in the different developmental stages of echinacea was determined.^[14]^ The information regarding the effect of the mycorrhizal inoculation on the biochemical composition of the different Echinacea species is not available in detail. Therefore, here we determined the changes in the biochemical composition of Echinacea as a result of mycorrhizal inoculation as they are environmentally friendly, economical and sustainable.

## METHODOLOGY

### Experimental setup and bioinoculation

The present experiment was conducted in the Department of Botany, Kurukshetra University, Kurukshetra in the open field condition from April 2020 to August 2020. An experiment was set up in a complete randomized block design (CRBD). Plants were grown on the moist filter paper and 10 days seedling was then transferred to the field containing soil:sand mixture whose physical and chemical composition is as follows.^[15,16]^

A soil culture of AMF, *Glomus mosseae* (having 82-86% colonized root pieces and 620-630 AM spores per 100 g) and *Gigaspora gigantean* (having 70-74% colonized root pieces and 500-520 AM spores per 100 g). Each AMF were then mass multiplied using sterile sand soil mixture (1:3) and Maize as host for 90 days, in greenhouse conditions. AMF are propagated as endomycorrhizal species as they are obligate symbiont. The native density of mycorrhizal spores in the experimental site was 24.5±7.09 per 10g soil, which was counted by the gridline intersect method.^[17]^ *Bacillus subtilis* (MTCC 1305) and *Pseudomonas fluorescens* (MTCC No. 103) were obtained from the Institute of Microbial Technology (Imtech), Chandigarh, India. Both of them were cultures in nutrient broth medium (NaCl, Peptone, Beef extract)and incubated at 32°C for 48 hrs to get a colony rate of 1×10^−11^ colony ml^−1^and 1×10^−9^ colony ml^−1^, respectively.

### Field preparation

First of all, the field of 7.5 × 10 feet was ploughed thoroughly for proper aeration and indigenous spores were counted before the planting of seedlings. Six flowerbeds of 1×1 m was made with a 10 cm alleyway as shown below. Fifteen + fifteen seedlings each of *E*. purpurea, *E*. *angustifolia*, *E*. *pallida* were then planted at 14 inches’ distance which was regularly watered by drip irrigation method. For each species, one flowerbed is kept as a control in which no inoculation was added. Still, other fields were subjected to consortium treatment as it was proven that consortium treatment enhances the growth and development of flowering plants.^[18,19]^

Inoculation contains 70-72% *G*. *mosseae* colonized maize roots (~1 cm) and 610-630 spores (per 100 g of maize rhizospheric soil), 65-67% *G*. *gigantea*colonized maize roots (~1 cm) and 580-600 spores (per 100 g of maize rhizospheric soil), *B*. *subtilis* and *P*. *fluorescence*. First bacterial inoculation was given during the planting of seedlings in the flowerbed by merely dipping the roots in the respective broth media for 10 min. After 10-13 days when plantlets affirm their roots (c.a. 10 mm), the second treatment of *G*. *mosseae* and *G*. *gigantea*was given by placing extra maize rhizospheric soil around the roots, to confirm the inoculation. Similarly, the second treatment of *B*. *subtilis* and *P*. *fluorescence* was given by respective sprinkling media around the roots.

### Parameters assessments

After 90 days of inoculum (DOI), out of fifteen plants from each plot, 10 plants were selected randomly for assessment. Morphological characters and biochemical as well as physiological parameters like whole plant dry weight, total chlorophyll, carotenoid, nitrogen, phosphorus, potassium, phenolics content, root essential oil, antioxidant activity and mycorrhization are assessed.Total dry weight was calculated by carefully uprooting the plant, weighing it and then oven-dry them at 55°C for 2 days, finally, by subtracting fresh weight with dry weight. Chlorophyll content was determined using optical absorbance at 620 and 940 nm. Whereas the total carotenoid content was determined by the Arnon’smethod.^[20]^. While total NPK content was calculated by the Bandyopadhyay *et al*. method.^[21]^. Ferric-Reducing Antioxidant was assessed by the modified method of Farhat *et al.*^[22]^ The method Wang *et al.*^[23]^ was used for determination of the total phenolic content. Whereas, Re *et al*. ^[24]^ method was used for the estimation of ABTS free radical-scavenging antioxidant activity. Total root essential oil was determined according tothehydro-distillation method of Mazza and Cottrell.^[25]^.

Mycorrhization started after 65-70 days of inoculation and was confirmed as well as quantified by Trypan blue staining.^[26]^. Further, the AM spore number and the percentage of root colonized was determined with the methods defined based on the method defined elsewhere.^[27]^

### Statistical analyses

Analysis of Variance (ANOVA) was conducted and one-way ANOVA was used to detect the differences among means of each treatment using SPSS (11.5 version) software package.^[28]^. The results of the experiment were analyzed for studying parameters between control and microbial-inoculated plants and the significance of differences was calculated using least significant differences (LSD) LSD (*P*≤0.05).

## RESULTS AND DISCUSSION

The findings of the experiment are about Biological traits; the bio-Physiological attributes mycorrhization pattern of three different species of echinacea is illustrated in [Table 1 to 2].

**Table 1:**
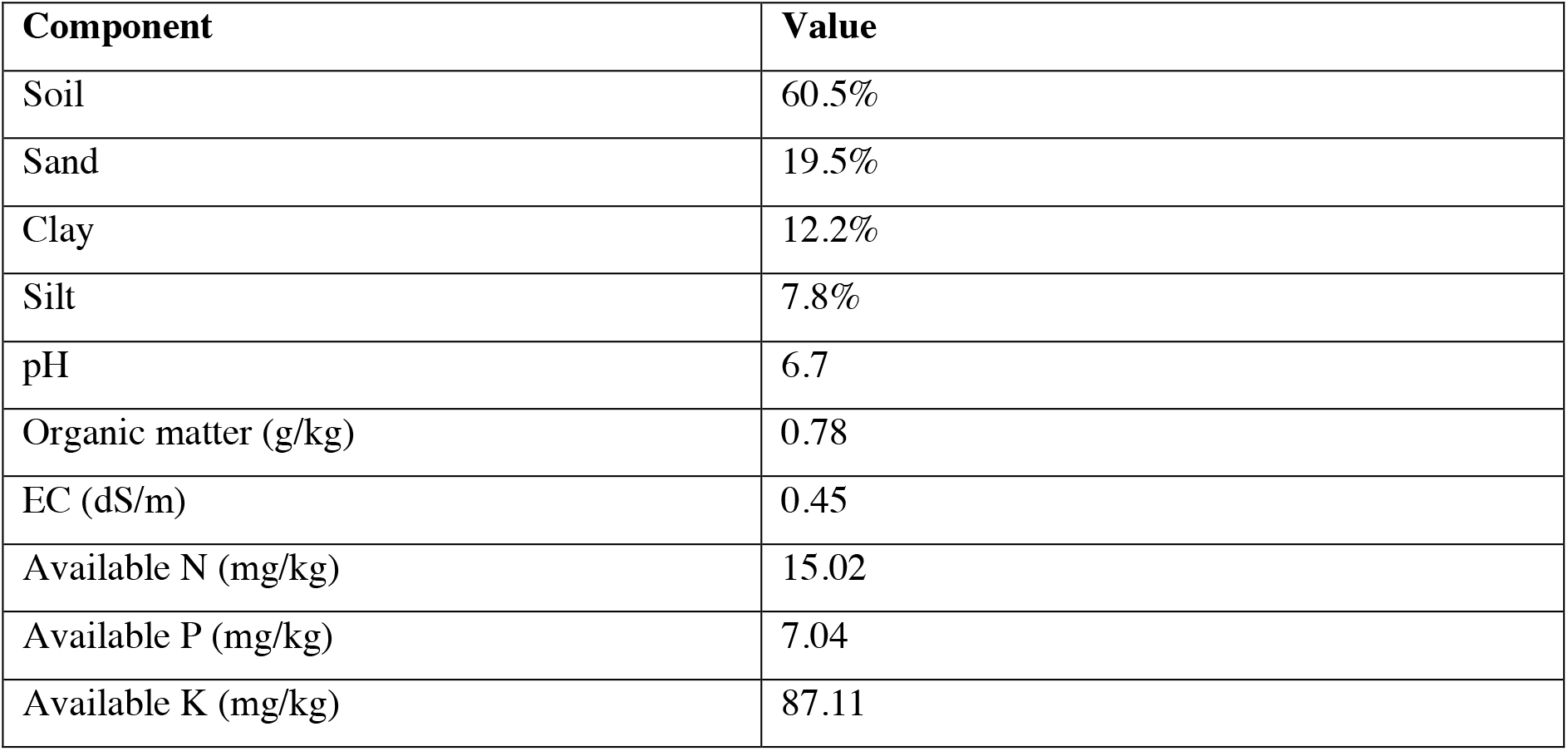
Composition of soil used for the cultivation of Echinacea species.

| Component | Value |
| --- | --- |
| Soil | 60.5% |
| Sand | 19.5% |
| Clay | 12.2% |
| Silt | 7.8% |
| pH | 6.7 |
| Organic matter (g/kg) | 0.78 |
| EC (dS/m) | 0.45 |
| Available N (mg/kg) | 15.02 |
| Available P (mg/kg) | 7.04 |
| Available K (mg/kg) | 87.11 |

**Table 1:** Changes in the total plant dry matter (g) and biochemical parameters of 3different echinacea specieswith/withoutmycorrhizal inoculation.

| Traits | Control | Consortium<br>(*Glomus mosseae*+*Gigasporagigantea*+*Pseudomonas fluorescens*<br>+ *Bacillus subtilis*) | ANOVA (2, 27) | LSD<br>($P \leq 0.05$) |
| --- | --- | --- | --- | --- |
| *Echinacea angustifolia* |  |  |  |  |
| Total plant dry matter (g) | 11.653±3.121<sup>b‡</sup> | 49.238 ±3.463<sup>a</sup> | 1.012.435 | 2.176 |
| Total chlorophyll (mg g<sup>-1</sup> FW) | 21.464±1.667<sup>b</sup> | 49.296±1.726<sup>a</sup> | 2.605.044 | 1.294 |
| Total carotenoid (mg g<sup>-1</sup> FW) | 11.045±0.778<sup>b</sup> | 25.645±0.895<sup>a</sup> | 3.580.926 | 0.574 |
| Total N content (%) | 1.882±0.264<sup>b</sup> | 2.583±0.344<sup>a</sup> | 319.115 | 0.201 |
| Total P content (ppm) | 0.304±0.113<sup>b</sup> | 0.778±0.123<sup>a</sup> | 132.912 | 0.091 |
| Total K content (ppm) | 0.941±0.289<sup>b</sup> | 1.799±0.151<sup>a</sup> | 240.418 | 0.155 |
| *Echinacea purpurea* |  |  |  |  |
| Total plant dry matter (g) | 10.753±2.458<sup>b‡</sup> | 45.738 ±3.878<sup>a</sup> | 787.304 | 2.432 |
| Total chlorophyll (mg g<sup>-1</sup> FW) | 23.793±3.609<sup>b</sup> | 52.796±3.818<sup>a</sup> | 732.364 | 2.783 |
| Total carotenoid (mg g<sup>-1</sup> FW) | 12.145±2.168<sup>b</sup> | 26.845±2.096<sup>a</sup> | 554.129 | 1.597 |
| Total N content (%) | 2.107±0.503<sup>b</sup> | 2.9631±0.554<sup>a</sup> | 51.816 | 0.396 |
| Total P content (ppm) | 0.305±0.097<sup>b</sup> | 0.887±0.095<sup>a</sup> | 225.479 | 0.072 |
| Total K content (ppm) | 0.934±0.292<sup>b</sup> | 1.839±0.439<sup>a</sup> | 27.366 | 0.279 |
| *Echinacea pallida* |  |  |  |  |
| Total plant dry matter (g) | 10.353±2.231<sup>b‡</sup> | 40.079±3.632<sup>a</sup> | 687.314 | 2.258 |
| Total chlorophyll (mg g<sup>-1</sup> FW) | 19.603±1.983<sup>b</sup> | 41.296±3.241<sup>a</sup> | 68.063 | 1.093 |
| Total carotenoid (mg g<sup>-1</sup> FW) | 9.249±0.889<sup>b</sup> | 23.707±0.855<sup>a</sup> | 2.602.873 | 0.653 |
| Total N content (%) | 0.939±0.305<sup>b</sup> | 1.947±0.364<sup>a</sup> | 42.460 | 0.251 |
| Total P content (ppm) | 0.243±0.101<sup>b</sup> | 0.697±0.098<sup>a</sup> | 218.137 | 0.075 |
| Total K content (ppm) | 0.622±0.269<sup>b</sup> | 1.424±0.335<sup>a</sup> | 26.115 | 0.243 |

±- Standard deviation; ‡- values in column followed by the same letter are not significantly different; $p \leq 0.05$ - LSD (least significant difference test); FW- Fresh Weight; ppm- Parts per million

**Table2:** Changes in thephysiological and mycorrhization characteristics of 3 differentEchinacea specieswith/withoutmycorrhizal inoculation.

| Traits | Control | Consortium<br>(*Glomus mosseae*+*Gigasporagigantea*+*Pseudomonas fluorescense*<br>+ *Bacillus subtilis*) | ANOVA (F2, 27) | LSD<br>($P \leq 0.05$) |
| --- | --- | --- | --- | --- |
| *Echinacea angustifolia* |  |  |  |  |
| Root essential oil (ml g<sup>-1</sup> FW) | 0.166±0.064<sup>b‡</sup> | 0.511±0.051<sup>a</sup> | 253.753 | 0.438 |
| ABTS Antioxidant (mm ml<sup>-1</sup>) | 15.343±1.711<sup>b</sup> | 23.164±1.425<sup>a</sup> | 791.887 | 1.214 |
| Ferric-Reducing Antioxidant (nmol μl<sup>-1</sup>) | 0.429±0.125<sup>b</sup> | 0.909±0.128<sup>a</sup> | 181.673 | 0.901 |
| Total phenolic (mEAG mL<sup>-1</sup>) | 15.383±1.043<sup>b</sup> | 21.644±0.896<sup>a</sup> | 1.799.164 | 0.707 |
| AM Spore No. | 38.600±8.275<sup>b</sup> | 221.300±8.069<sup>a</sup> | 552.277 | 4.754 |
| AM Root Colonization % | 21.400±6.186<sup>b</sup> | 74.200±6.442<sup>a</sup> | 2.758.162 | 6.576 |
| *Echinacea purpurea* |  |  |  |  |
| Root essential oil (ml g<sup>-1</sup> FW) | 0.1852±0.093<sup>b‡</sup> | 0.613±0.123<sup>a</sup> | 207.758 | 0.082 |
| ABTS Antioxidant (mm ml<sup>-1</sup>) | 14.903±2.061<sup>b</sup> | 24.982±2.129<sup>a</sup> | 495.176 | 1.571 |
| Ferric-Reducing Antioxidant (nmol μl<sup>-1</sup>) | 14.033±2.494<sup>b</sup> | 0.999±0.174<sup>a</sup> | 426.688 | 1.631 |
| Total phenolic (mEAG mL<sup>-1</sup>) | 0.504±0.092<sup>b</sup> | 24.167±1.805<sup>a</sup> | 62.778 | 0.104 |
| AM Spore No. | 37.600±8.154<sup>b</sup> | 231.300±9.849<sup>a</sup> | 997.800 | 11.368 |
| AM Root Colonization % | 22.4000±4.971<sup>b</sup> | 78.700±5.618<sup>a</sup> | 858.682 | 3.974 |
| *Echinacea pallida* |  |  |  |  |
| Root essential oil (ml g<sup>-1</sup> FW) | 0.153±0.068<sup>b‡</sup> | 0.4711±0.064<sup>a</sup> | 613.859 | 0.051 |
| ABTS Antioxidant (mm ml<sup>-1</sup>) | 13.603±1.719<sup>b</sup> | 22.055±1.778<sup>a</sup> | 550.412 | 1.310 |
| Ferric-Reducing Antioxidant (nmol μl<sup>-1</sup>) | 13.528±1.114<sup>b</sup> | 0.851±0.125<sup>a</sup> | 984.040 | 0.884 |
| Total phenolic (mEAG mL<sup>-1</sup>) | 0.339±0.137<sup>b</sup> | 19.765±1.241<sup>a</sup> | 104.020 | 0.098 |
| AM Spore No. | 33.900±5.665<sup>b</sup> | 195.200±7.871<sup>a</sup> | 3.445.503 | 5.138 |
| AM Root Colonization % | 19.600±4.501<sup>b</sup> | 69.600±3.949<sup>a</sup> | 1.052.777 | 3.179 |

±- Standard deviation; ‡- values in column followed by the same letter are not significantly different;
AM- Arbuscular Mycorrhizal $p \leq 0.05$ - LSD (least significant difference test); FW- Fresh Weight; mEAG $\text{mL}^{-1}$ - Equivalent mg of gallic acid per milliliter of extract
mm $\text{mL}^{-1}$ - millimolar per milliliter of extract (=Trolox Equivalent Antioxidant Capacity)
ABTS-2,2'-azino-bis (3-ethylbenzothiazoline-6-sulfonic acid); $\text{nmol } \mu\text{L}^{-1} = \text{mM Fe}^{2+}$ equivalents.

## Biological traits

In this experiment, biological traits, total plant dry matter, total chlorophyll, total carotenoid, total N content, total P content, and total K content were found increased over control [Table 1]. The consortium showed a significant effect on the biological traits of Echinacea spp. The highest plant dry matter was higher under *E. angustifolia* (49.29 g) followed by *E.purpurea and E. pallida*[Table 1]. Total chlorophyll content was higher under *E. purpurea* (52.79 mg g^−1^) than the *E. angustifolia* (49.29 mg g^−1^)and *E. pallida* (41.29 mg g^−1^).Total carotenoid was higher under *E. angustifolia* (25.64 mg g^−1^) followed by *E. purpurea* (26.84 mg g^−1^) and minimum Cartenoid was found under *E.pallida (*23.70 mg g^−1^) [Table 1]. Total N content was found higher under consortium over control. The highest N content was under *E. purpurea* (2.96%) followed by *E. angustifolia* (2.58%) and *E. apallia*(1.94%) [Table 1]. Total P and K content were significantly higher under consortium than the control plant [Table 2]. The highest P was higher under *E. purpurea* (0.88 ppm) followed by *E.angustifolia* (0.77 ppm) and *E. pallida* has lowest P content[Table 1]. The same result was found for K content in echinacea species over the control treatment. Highest K was found under *E. purpurea* (1.83 ppm) followed by *E. angustifolia* (1.79 ppm) and lowest under *E.pallida* (1.42 ppm) [Table 1].

## Physiological and mycorrhization characteristics

Biophysical traits (Root essential oil, ABTS Antioxidant, ferric- reducing antioxidant and total phenolic) and mycorrhization pattern (AM spore, AM root colonization) were higher under consortium than the control plant [Table 2]. Root essential oil was higher under *E. purpurea* (0.61 ml g^−1^) followed by *E. angustifolia* (0.511ml g^−1^) and lowest in *E. pallia* (0.47 ml g^−1^) [Table 2]. From Table 1 it is clear that biophysical attributes were higher under consortium than the control plant and AM spore no. and Am root colonization were also higher under mycorrhizal treatment than the control [Table 2]. Among echinacea species. biophysical traits and mycorrhization pattern were higher in *E. purpurea* followed by*E. Angustifolia* lowest under *E. pallida* [Table 2]). ABTS antioxidant was highest under *E. purpurea* (24.98 mm ml^−1^) and followed by *E. angustifolia* and *E. purpurea* (23.16 mm ml^−1^) [Table 2]. *E. purpurea* showed the highest ferric- reducing antioxidant (0.99 nmol μl–1) and total phenolic (24.16 mEAG mL^−1^) [Table 2]. *E. angustifolia* and *E. pallida* showed ferric- reducing antioxidants 0.90 nmol μl–1, 0.85nmol μl–1 respectively. The total phenolic compound was lowest under *E. pallida* (19.76mEAG mL^−1^) [Table 2].

The differences between the AM spore no., AM root colonization (%) are illustrated in [Table 2]. Both the parameters werefound significantly higher under consortium than the control. AM spore no. was found highest in *E. purpurea* than the other species of echinacea [Table 2].AM spore no. *E. angustifolia* and *E. purpurea*were221.30and 231.30 respectively. AM root colonization is also higher under *E. purpurea*than the other echinacea species. AM root colonization was higher under *E. purpurea* depicted in [Table 2]. *E*. *angustifolia* and *E. pallida* showed AM root colonization 74.2%and 69.6%, respectively [Table 2].

After 90 days of inoculum, all the plants having bioinoculants showed better results as compared to control. According to the results, incorporating mycorrhizae in echinacea species. the biochemical composition was increased. AMF was very significant for improvising the biochemical composition. Applying plant growth-promoting rhizobacteria (PGPR) and mycorrhizal fungi improve the traits and yield in *E*. *purpurea*. The mixture of bacteria with mycorrhizal inoculum and shoots was treated with biofertilizers to enhance the biochemical composition.^[29]^ The phenolic compound was higher in *E. purpurea* followed by *E. angustifolia* and *E. pallida.* ^[30]^ There was variability among *echinacea* species. for biochemical composition. Variable positive effects of mycorrhiza were found for different varieties of echinacea. Similar results were found in root colonization by mycorrhizal fungi with *E. purpurea*. Increasing root colonization with the help of biofertilizers helps to improve the phenolic compound of plant roots such as cynarin, cichoric and caftaric acid.^[31]^ Soil fertilization with the biochemical fertilizers such as AMF and *Pseudomonas* species have improved growth and yield components.^[32–36]^ AMF and PFB were more useful for increasing plant nutrients, improving leaf relative water content and decreased ion leakage.^[37]^ *E. angustifolia* showed increased echinacoside content by applying K. In all addition to increasing absorption of nutrients, development by plant hormones, controlling plant pathogens and some other factors.^[38,39]^ AMF improved the host plants in many ways such as the uptake of phosphorus and other nutrients, increased plant growth, plant height, leaf area, fresh\dry weight of shoots and roots.^[40,41]^

## CONCLUSION

From the present experimentation, it can be concluded that bioinoculants modify the phytohormones, water uptake efficiency and photosynthetic activity. This results in better plant growth and increased bio-physico chemical parameters. By this, we can draw attention toward the commercialization of plant products, mostly medicinal plants and their products, as this microbial association is sustainable and beneficial, not only for plants but for the soil ecosystem too. Medicinal values of any plant can be altered, this can also increase the economy of the country and even application of chemical fertilizers can be reduced. The result of mycorrhizal association was very effective for plant growth and increased bio-physicochemical properties than the control plants. Out of the current tests, it might be recognized that the phytohormones are changed by bio inoculants, water uptake success in addition to photosynthetic activity. This produces better place growth and improved bio physiochemical parameters. The results of the mycorrhizal association were significant for plant growth plus improved biophysiochemical characteristics when compared with the management plants.

